# Pericyte dysfunction alone may be insufficient to drive tick-borne orthoflavivirus NS1-mediated microvascular permeability

**DOI:** 10.64898/2026.01.29.702573

**Authors:** Emma Brown, Justin Duruanyanwu, Paola Campagnolo, Kevin Maringer

**Affiliations:** The Pirbright Institute, Ash Road, Pirbright, Surrey, GU24 0NF, United Kingdom; School of Biosciences, University of Surrey, Guildford, Surrey, GU2 7XH. United Kingdom

**Author notes:** To whom correspondence should be addressed: Paola Campagnolo, Kevin Maringer.

**Keywords:** flavivirus NS1, Alkhumra (Alkhurma) haemorrhagic fever virus (AHFV), Kyasanur forest disease virus (KFDV), microvascular leakage, endothelial cells, perivascular cells (pericytes)

## Abstract

The emerging tick-borne flaviviruses Alkhumra haemorrhagic fever virus (AHFV) and Kyasanur Forest disease virus (KFDV) can cause severe haemorrhagic disease, yet the mechanisms driving this vascular leakage remain unclear. The microvascular capillaries and post-capillary venules affected during orthoflavivirus-related haemorrhagic disease consist of an endothelial cell barrier ensheathed in and supported by perivascular cells (pericytes), and we recently identified a critical role for pericytes in amplifying dengue microvascular dysfunction. The orthoflavivirus non-structural protein 1 (NS1) has been implicated in endothelial dysfunction and vascular leakage for several tick- and mosquito-borne flaviviruses. Here, we examined whether NS1 from AHFV and KFDV disrupts microvascular barrier integrity in pericyte-endothelial cell cocultures. We found AHFV and KFDV NS1 to disrupt the ability of pericytes to support endothelial cell function, which is required for maintenance of the microvascular barrier. However, under our experimental conditions, we detected no endothelial hyperpermeability upon NS1 treatment in endothelial cells cultured alone or in combination with pericytes. Our findings suggest that the concomitant impairment of both endothelial cell and pericyte function may be required for NS1 to induce microvascular hyperpermeability. Our work highlights the potentially synergistic effects of microvascular cells during orthoflavivirus haemorrhage and emphasises the need for further work into the mechanisms of vascular dysfunction during AHFV and KFDV infection.

## Introduction

Over the past five decades, arboviral epidemics, particularly those caused by viruses in the *Flaviviridae* family, have (re-)emerged as significant public health threats. Notable examples include the 2015 Zika virus (ZIKV) epidemic in Latin America, associated with neurological complications and congenital abnormalities; the spread of West Nile virus (WNV) across the Americas and now Europe; and increasingly common and severe dengue virus (DENV) outbreaks in tropical and subtropical regions [1]. Two recently emerging tick-borne orthoflaviviruses are Alkhumra haemorrhagic fever virus (AHFV) and Kyasanur Forest disease virus (KFDV). AHFV was first identified in Saudi Arabia in 1995 and has since caused 611 reported cases, though this is likely an underestimate, as only hospitalized patients are reported and diagnosed [2–4]. KFDV, first detected in Karnataka, India in 1957, now causes around 400–500 cases annually, primarily along India’s western coast [5]. Both viruses cause flu-like febrile symptoms with or without haemorrhagic manifestations, including epistaxis, purpura, bleeding gums, gastrointestinal bleeding, and menorrhagia, as well as neurological complications [4,6–8]. The proportion of patients presenting with haemorrhagic symptoms ranges from 20–55%, with estimated case fatality rates of up to 25% for AHFV and 3–5% for KFDV [6]. Despite the clinical severity, how vascular leakage arises during these infections remains poorly understood.

Orthoflavivirus-induced haemorrhage has multifactorial causes, but the secreted non-structural protein 1 (NS1) can directly initiate endothelial dysfunction and vascular leakage [9–13]. NS1 has been shown to impair endothelial function by degrading components of the endothelial glycocalyx and disrupting junction proteins essential for cell–cell adhesion, among other mechanisms [14–17]. Beyond its direct effects on endothelial cells, we recently demonstrated that NS1-induced hyperpermeability is amplified in the presence of perivascular cells (pericytes) [18]. Pericytes are mesenchymal-like cells that envelop endothelial cells in the microvascular capillaries and post-capillary venules affected during orthoflavivirus haemorrhage, playing a critical role in maintaining vascular stability through both direct physical contact with endothelial cells and reciprocal paracrine signalling [18,19]. These findings suggest that NS1 disrupts not only endothelial barrier integrity but also the coordinated intracellular interactions required for microvascular homeostasis.

In this study, we investigated whether NS1 from AHFV and KFDV can induce microvascular dysfunction similarly to other orthoflaviviruses. Interestingly, although NS1 from both viruses disrupted the ability of pericytes to support endothelial cell function in an angiogenesis assay, in addition to other physiological impacts on pericytes, no impact on endothelial permeability was observed in endothelial cell monocultures or pericyte-endothelial cell cocultures in the cell models tested here. Our data demonstrate that while pericytes amplify NS1-induced endothelial dysfunction, pericyte dysfunction alone may be insufficient to cause orthoflavivirus microvascular leakage in the absence of endothelial cell effects, highlighting potentially synergistic impacts on microvascular cells during orthoflavivirus haemorrhage.

## Materials and Methods

### Cell culture

Human umbilical vein endothelial cells (HUVECs) were obtained from PromoCell (Heidelberg, Germany); liver sinusoidal endothelial cells (LSECs) from Innoprot (Derio, Spain), and human hepatic stellate cells (HSCs) from Zen-Bio (Durham, NC USA). HUVECs, LSECs and HSCs were grown to 90% confluency in endothelial growth media supplemented with specific growth factors (PromoCell for HUVECs; Innoprot for LSECs) or Human Stellate growth medium (HSCs, Zen-Bio) in culture plates precoated with 15 µg/mL bovine fibronectin (LSECs) or 50 µg/mL rat tail collagen I (HSCs). HUVECs and LSECs were used between passages 3–8; HSCs between passages 3–5. Chinese Hamster Ovary (CHO) DG44 cells were kindly provided by Prof John McVey (University of Surrey), maintained in suspension in CD DG44 media (Gibco) supplemented with 8 mM L-Glutamine and 1.8% Pluronic F-68 (Gibco, ThermoFisher, Waltham, MA, USA) at 37°C with 5% CO₂ on a shaker at 120 rpm. The cell culture media was changed twice a week, and the cells were split 1:2 when they reached 1.5 × 10⁶ cells/mL.

### NS1 sequence analyses

All of the available 22 AHFV and 47 KFDV NS1 amino acid sequences were retrieved from the Virus Pathogen Resources (ViPR) database (bv-brc.org) [20,21] in FASTA format (access date 07/05/2026). Overall sequence conservation was determined using BLASTp with standard parameters (blast.ncbi.nlm.nih.gov) [22]. Residue-by-residue sequence diversity analysis was performed by aligning all retrieved NS1 sequences for AHFV and KFDV separately to identify amino acid differences at each position of the NS1 protein sequence. Multiple sequence alignment was carried out using Molecular Evolutionary Genetics Analysis (MEGA) version 12.1.2 employing the ClustalW algorithm with default parameters [23,24]. Consensus sequences were generated from the multiple sequence alignments using the EMBOSS Cons tool implemented through the EMBL-EBI Job Dispatcher framework (EMBL-EBI, access date 08/06/2026). Consensus residues were assigned according to the most frequent amino acid at each alignment position. Consensus sequences were subsequently aligned with the NS1 sequences used to generate recombinant AHFV and KFDV NS1 using MEGA version 12.1.2.

### Production of recombinant soluble NS1

Plasmids encoding Streptavidin-tagged NS1 from AHFV (pKM74; viral reference sequence NCBI accession JF416957.1) or KFDV (pKM79 accession JF416958.1) were transfected into CHO DG44 cells using FreeStyle MAX reagent (ThermoFisher), followed by selection using 0.3 µg/mL geneticin (G418 sulphate, ThermoFisher). Cell culture supernatant containing NS1 was harvested once cell density exceeded 1.8 × 10⁶ cells/mL with viability above 85%. Strep-tagged NS1 proteins were purified using Strep-Tactin Sepharose gravity flow columns (IBA Lifesciences, Göttingen, Germany). Purified protein eluates were pooled and quantified with a Qubit fluorometer (ThermoFisher). Proteins were stored at –80°C in storage buffer (2.5% trehalose, 100 mM arginine, and 0.2% ultrapure bovine serum albumin) to prevent degradation. To confirm that purified NS1 was in the expected tetrameric or hexameric form [25], purified NS1 was passed through a 100 kDa Amicon Ultra size separation column (Merck, Darmstadt, Germany) and analysed by SDS-PAGE and western blot. Plasmids have been deposited with Addgene (addgene.org, Watertown, MA USA) with Addgene ID 250163 (pKM74, AHFV) and 250164 (pKM79, KFDV). NS1 preparations were not tested for residual host cell proteins or other contaminants; however, an eluate control (protein purification eluates from cells not expressing NS1; negative control) was included in all experiments and would be expected to contain the same contaminants. His-tagged DENV-2 NS1 and DENV-2 N207Q NS1 were purchased from The Native Antigen Company (Oxford, UK).

### Angiogenesis assays

Each well of a 96-well plate was coated with 45 μl of basement membrane matrix (Geltrex®, Gibco) on ice and allowed to solidify at 37 °C for 30 min. Next, single cultures of 2 × 10^4^ cells/well of HUVECs or cocultures of 2 × 10^4^ cells/well of HUVECs and 5 × 10^3^ cells/well of HSCs were seeded on top of the Geltrex® and immediately treated with a single dose of 1 µg/mL NS1 (final concentration) or an eluate control (negative control). Cells were incubated for 6 h in a 37 °C incubator with 5% CO_2,_ before two bright-field images were taken from each well at a magnification of 10X. Images were quantified using Fiji software (version 2.1.0) with the CMP vascular analyser macro plugin (version 2), which automatically quantified the number of segments, number of junctions and total branching length [26]. Segment width was quantified manually as described previously [9].

### Viability assays

HSCs were seeded in 96-well plates at 2 × 10³ cells per well and incubated for 24 h prior to treatment. Cells were then treated with NS1 (1 µg/mL) for 24 h in triplicate. Cell viability was assessed using an MTT assay (Cayman Chemicals Company, Ann Arbor, MI USA) with MTT (500 µg/mL) added for 4 h followed by solubilisation of formazan crystals using MTT solvent with shaking (100 rpm, 15 min). Plates were incubated overnight before absorbance was measured at 570 nm using a SpectraMax ID3 microplate reader (Molecular Devices, LLC, San Jose, CA USA). Relative absorbance was calculated by subtracting background values from blank wells and normalising to untreated controls.

### RT-qPCR assays

HSCs were seeded at 5 × 10⁴ cells per well in six-well plates (Sarstedt, Nümbrecht, Germany) and treated when sub-confluent with 1 μg/mL NS1 for 24 h. Cells were lysed in TRI Reagent (Merck) and RNA extracted using the Zymo RNA Microprep Kit (Cambridge Bioscience Ltd, Cambridge, UK) according to the manufacturer’s instructions. RNA concentration was measured using a DeNovix spectrophotometer (DeNovix Inc., Wilmington, DE USA).

For cDNA synthesis, 100 ng RNA was reverse-transcribed using the RevertAid First Strand cDNA Synthesis Kit (ThermoFisher) according to the manufacturer’s instructions. qPCR was performed using PowerUp SYBR Green Master Mix (ThermoFisher) in 96-well MicroAmp plates (ThermoFisher) with gene-specific primers (1 μM final concentration) targeting angiopoietin-1 (Ang-1) (NCBI Gene ID: 284, Merck), VEGF (NCBI Gene ID: 7422, Merck), interleukin 6 (IL-6) (NCBI Gene ID: 3569, Merck) and β-actin (internal control; NCBI Gene ID: 60, Merck). Primer sequences were as follows: Ang-1 forward 5’- AGAACCTTCAAGGCTTGGTTAC-3’ and reverse 5’- GGTGGTAGCTCTGTTTAATTGCT- 3’; VEGF forward 5′-TGCAGATTATGCGGATCAAACC-3′ and reverse 5′- TGCATTCACATTTGTTGTGCTCTAG-3′; IL-6 forward 5’- AATGCCAGCCTGCTGACGAA-3’ and reverse 5’-CTGAGGTGCCCATGCTACAT-3’; β- actin forward 5’-AGAGCTACGAGCTGCCTGAC-3’ and reverse 5’- AGCACTGTGTTGGCGTACAG-3’. cDNA was added to qPCR reactions at a final dilution of 1:5. Plates were run on a QuantStudio 5 system using QuantStudio Real-Time PCR Software version 1.7.2 under the following cycling conditions: 50 °C for 2 min, 95 °C for 10 min, followed by 40 cycles of 95 °C for 15 s and 60 °C for 1 min. Relative gene expression was calculated using the 2−ΔΔCt method, with Ct values normalised to β-actin and expressed relative to the untreated control.

### Microvascular permeability assays

For the transendothelial electrical resistance (TEER) assay using the Electric Cell-substrate Electrical Impedance Sensing (ECIS) Z-Theta system (Applied Biophysics, Troy, NY USA), 8W10E+ arrays were pre-treated overnight at 37 °C with 400 µL of culture media prior to coating with 0.5 µg/mL human fibronectin (Merck) for HUVECs or 15 µg/mL bovine fibronectin (Innoprot) for LSECs. Cells were then seeded at 1.8 × 10⁵ cells/well in 400 µL media per well and incubated for 24 h. Monolayer confluence was confirmed by stable resistance readings >1000 Ω at 4 kHz before treatment with a final concentration of 1 µg/mL, eluate control, or 100 ng/mL tumour necrosis factor alpha (TNF-α) (PeproTech, Cranbury, NJ USA) for HUVEC and LSEC monocultures. Alternatively, endothelial cell monocultures were treated for 24 h with pre-conditioned media from 1 × 10⁵ cells/well HSCs. HSCs were cultured in 6-well plates and, upon reaching confluence, treated for 24 h with 1 µg/mL NS1 or 100 ng/mL TNF-α in endothelial growth medium before the collection of the cell supernatant. Resistance readings were recorded every two hours and normalised to each sample prior to treatment.

For coculture TEER assay using the EVOM system (World Precision Instruments, Sarasota, FL USA), HSCs were seeded on the underside of 0.4 µm pore-size 24-well plate Transwell inserts (Greiner Bio-One Ltd, Gloucestershire, UK) at 2.4 × 10⁴ cells/well in HSC media and incubated for 4 h. Inserts were flipped upright into 24-well plates and cultured for an additional 48 h in HSC media. Before seeding endothelial cells, the apical side of the insert was coated with either 0.5 µg/mL human fibronectin (HUVECs) for 20 min at room temperature or 15 µg/mL bovine fibronectin (LSECs) overnight at 37 °C. HUVECs or LSECs were seeded at 8 × 10⁴ cells/well in 200 µL and allowed to form monolayers over 48 h in endothelial growth media. Endothelial cell monocultures were similarly prepared but without the addition of HSCs. Cells were treated with a final concentration of 1 µg/mL or 5 µg/ml NS1, eluate control, or 100 ng/mL TNF-α. TEER was measured at times indicated in figures using an EVOM volt-ohm meter (World Precision Instruments). Absolute TEER was calculated by subtracting the blank (cell-free insert) and multiplying by the surface area of the insert. Relative TEER values were normalized to untreated endothelial cell monocultures at each corresponding time point. After 24 h of NS1 treatment, permeability was also assessed using a dextran permeability assay by introducing a final concentration of 250 µg/mL Cascade Blue dextran (10,000 MW; ThermoFisher) into the apical side of the transwell insert. After 1 h, 100 µL of culture media was collected from the basolateral chamber and fluorescence measured on a SpectraMax iD3 plate reader at excitation/emission: 400/450 nm (Molecular Devices). Media-only controls were used to determine background fluorescence.

### Images and Statistical Analysis

Graphs were plotted in R (version 4.4.0; R Core Team, 2025) using RStudio (version 2022.2.2.485; Posit, 2025) with *ggplot2* [27], *scales* [28] and *cowplot* [29] packages for visualisation. Individual data points were plotted, with data shown as mean ± standard error of the mean (SEM) for bar graphs, or mean ± SEM for line graphs. Experiments were performed with three biological replicates, each comprising three or four technical replicates. Statistical analysis was performed in R (version 4.4.0) within RStudio (2022.2.2.485) using the *lme4* [30], *MASS* [31], *multcomp* [32], and *XLConnect* [33] packages. Statistical comparisons represent all pairwise differences between treatment groups at each time point and were analysed using a linear mixed-effects model with experiment included as a random intercept. Multiple comparisons were performed using Tukey-adjusted post-hoc tests. Technical replicates were treated as repeated measurements within each biological experiment and were not considered independent experimental units. Differences were considered statistically significant at P < 0.05.

## Results

### Sequence comparison, cloning and purification of recombinant AHFV and KFDV NS1

To gain a better understanding of NS1 sequence variation, we first compared all 22 AHFV and all 47 KFDV NS1 amino acid sequences available in the Virus Pathogen Resources (ViPR) database (bv-brc.org) [20,21]. The currently available AHFV and KFDV NS1 amino acid sequences exhibit an overall sequence identity of 98.87% within each of these viral species, and between the two viruses the minimum NS1 amino acid sequence identity is 94.33%. The minimum sequence identity at any particular position within the NS1 protein sequence is 95.2% for AHFV and 94.92% for KFDV, which is observed for a minority of residues (Figure 1A).

**Figure 1.**
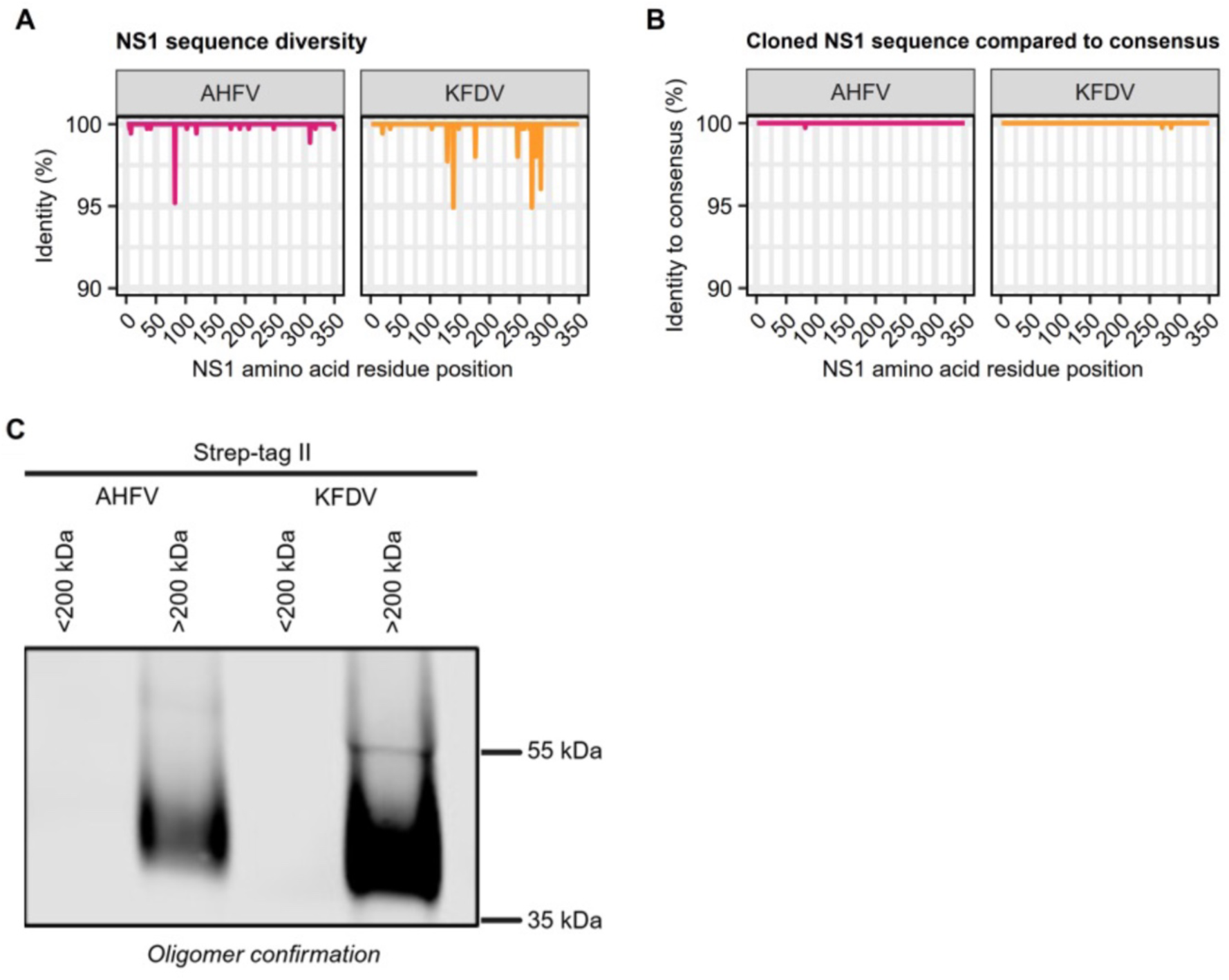
Production and characterisation of oligomeric AHFV and KFDV NS1 proteins. (A) Sequence diversity across all AHFV and KFDV NS1 sequences available in the ViPR database (22 for AHFV and 47 for KFDV). (B) Comparison of the cloned AHFV and KFDV NS1 sequences used in this study with consensus NS1 sequences generated from all available sequences in the ViPR database. (C) Purified NS1 proteins were separated using an Amicon ultrafiltration device with a 200 kDa molecular weight cut-off to distinguish dimeric (<200 kDa) from higher-order oligomeric (>200 kDa) forms. SDS-PAGE analysis confirmed the presence of both AHFV and KFDV NS1 in the oligomer-containing fractions.

We selected and cloned NS1 sequences from AHFV (NCBI accession JF416957.1) and KFDV (JF416958.1) that differed from the consensus by only one or two residues respectively at the most variable position for each virus (Figure 1B). None of the residues in question have been reported to play a role in NS1-mediated permeability based on the current literature [34]. These particular NS1 sequences were chosen because the AHFV sequence is one of the most complete AHFV genome sequences available, and because this particular KFDV isolate has previously been shown to cause systemic disease and a cytokine storm in experimentally infected macaques [35]. Stable CHO cell lines expressing AHFV and KFDV NS1 were generated and NS1 was affinity purified from the culture supernatant. To confirm that our purified recombinant NS1 proteins formed higher order oligomers, size exclusion columns were used to separate purified NS1 into its dimeric form (<200 kDa) and higher-order oligomeric forms (>200 kDa). Both AHFV and KFDV NS1 were thus confirmed to have been purified in their oligomeric states by western blot (Figure 1C). Taken together, these data validated the suitability of the NS1 proteins used in this study for downstream functional assays.

### AHFV and KFDV NS1s impair the ability of pericytes to support endothelial cell function

We previously showed that DENV-2 NS1 impairs the ability of pericytes to support endothelial cell function, which can be measured in a 3D angiogenesis assay in which endothelial cells form a network of vessel-like structures that becomes more extensive and complex in the presence of pericytes [9]. To investigate whether AHFV and KFDV NS1s similarly interfere with pericyte-endothelial cell interaction, human umbilical vein endothelial cells (HUVECs) and hepatic stellate cells (HSCs, liver pericytes) were cocultured on a basement membrane and treated with 1 µg/mL AHFV or KFDV NS1. HSCs were selected for these experiments as they are widely regarded as the liver-resident pericytes of the sinusoidal microvasculature and play a key role in regulating endothelial stability and vascular permeability [18]. Liver pericytes were chosen based on their relevance to AHFV and KFDV pathogenesis [35,36]. As observed previously [9], the presence of pericytes visibly increased the complexity of the vessel-like network (Figure 2A). Treatment with all NS1s tested, but not a control eluate generated by the same method as AHFV and KFDV NS1, visibly reduced this ability of pericytes to support the formation of the endothelial network relative to the untreated cocultures (Figure 2A). When these differences were quantified (quantification approach shown in Figure 2B), we observed a statistically significant decrease in all indicated measures of the overall complexity of the vessel-like network following treatment with DENV-2 NS1 (positive control), AHFV NS1 or KFDV NS1 compared to the untreated coculture control (Figure 2C-F). Treatment with the eluate control did not significantly reduce the number of segments, total branching length, nor the width of the segments, but did reduce the number of junctions significantly to a lesser degree than the NS1 treatments. This may be due to salts or other minor contaminants carried over from our recombinant protein expression and purification workflow, and emphasises the need for this control in addition to untreated cells. Overall, these data confirm that, as was previously shown for DENV-2 NS1, AHFV and KFDV NS1 disrupt the ability of pericytes to support endothelial cell functionality.

**Figure 2:**
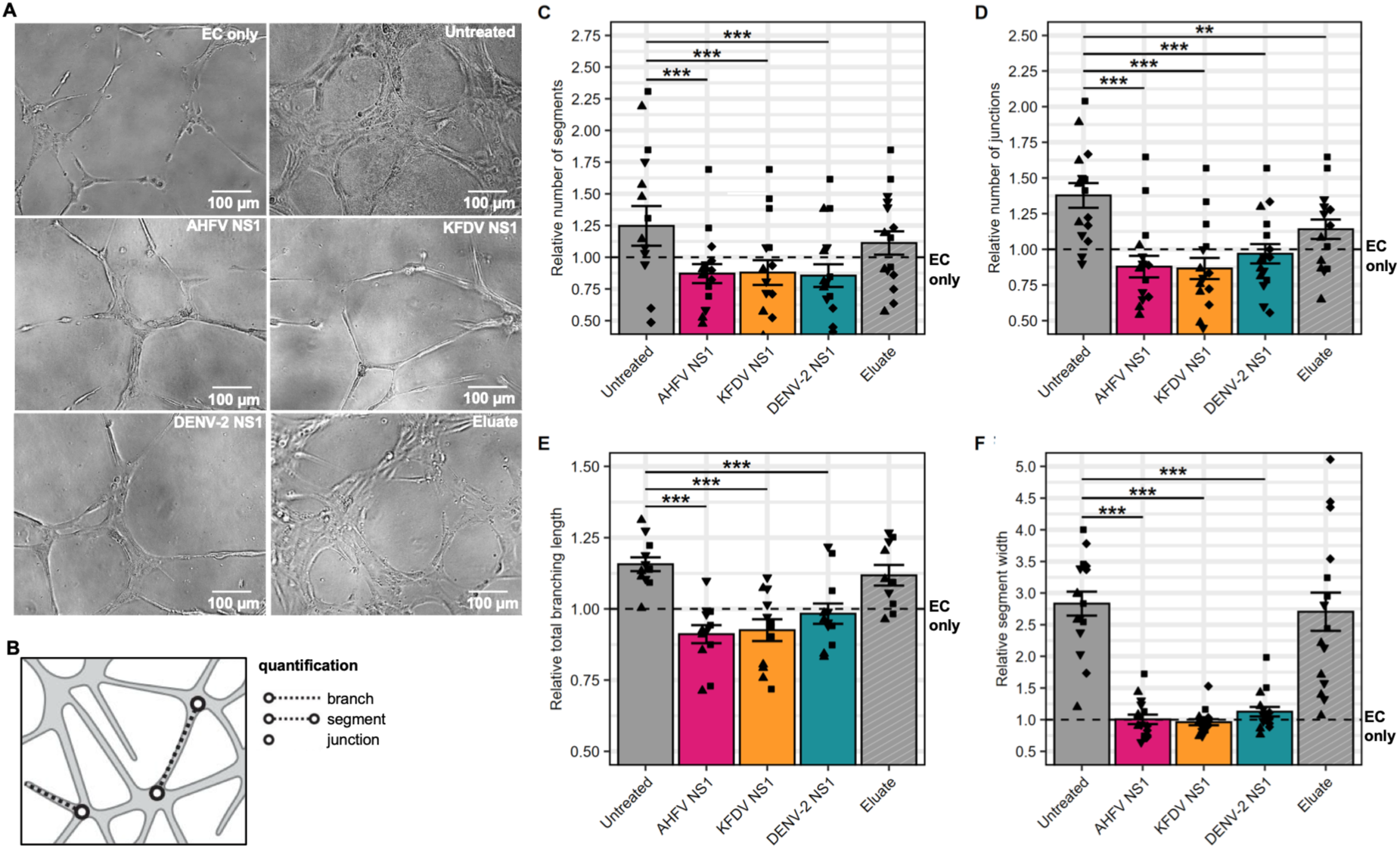
AHFV and KFDV NS1s disrupt pericytes’ ability to support endothelial cell vessel-like structures in a 3D angiogenesis model. (A) Representative images of HUVECs forming vessel-like structures in 3D Geltrex® matrix in the presence of HSCs, with or without 1 µg/mL NS1 treatment. Scale bars, 100 µm. (B) Angiogenesis assay quantification method. “Junctions” connect vessel-like structures; “branches” connect to junctions at one end; “segments” connect at both ends. “Total branching length” includes all branches; counts reflect the total number of segments, branches and junctions within field of view. (C-F) Quantification of vessel-like structures in pericyte-endothelial cell cocultures following NS1 treatment, normalised to untreated HUVECs (dashed line). Data points represent technical replicates within an experiment with different symbol shapes corresponding to different experiments (N=4, n=4 for the angiogenesis assays and N=3, n=2 for the PCR assays). Bars represent the mean, and error bars represent SEM. Statistical comparisons represent all pairwise differences between treatment groups. Indicated significance is for treatment versus untreated cocultures (Untreated); * *P* < 0.05; ** *P* < 0.01; *** *P* < 0.001.

### AHFV and KFDV modulate pericyte physiology in a number of ways

To further investigate the impact of NS1 on pericytes, we assessed whether NS1 treatment affected the viability of HSCs (Figure 3A) as well as canonical pericyte signalling pathways known to regulate microvascular integrity (Figure 3B–D). Similar to our previous data with DENV NS1 [9], cell viability was not significantly affected following treatment with AHFV NS1 compared to untreated HSCs (Figure 3A). In contrast, KFDV NS1 significantly decreased HSC viability (Figure 3A). These findings indicate that AHFV NS1 is not cytotoxic to pericytes under the conditions tested, whereas KFDV NS1 did reduce HSC viability.

**Figure 3:**
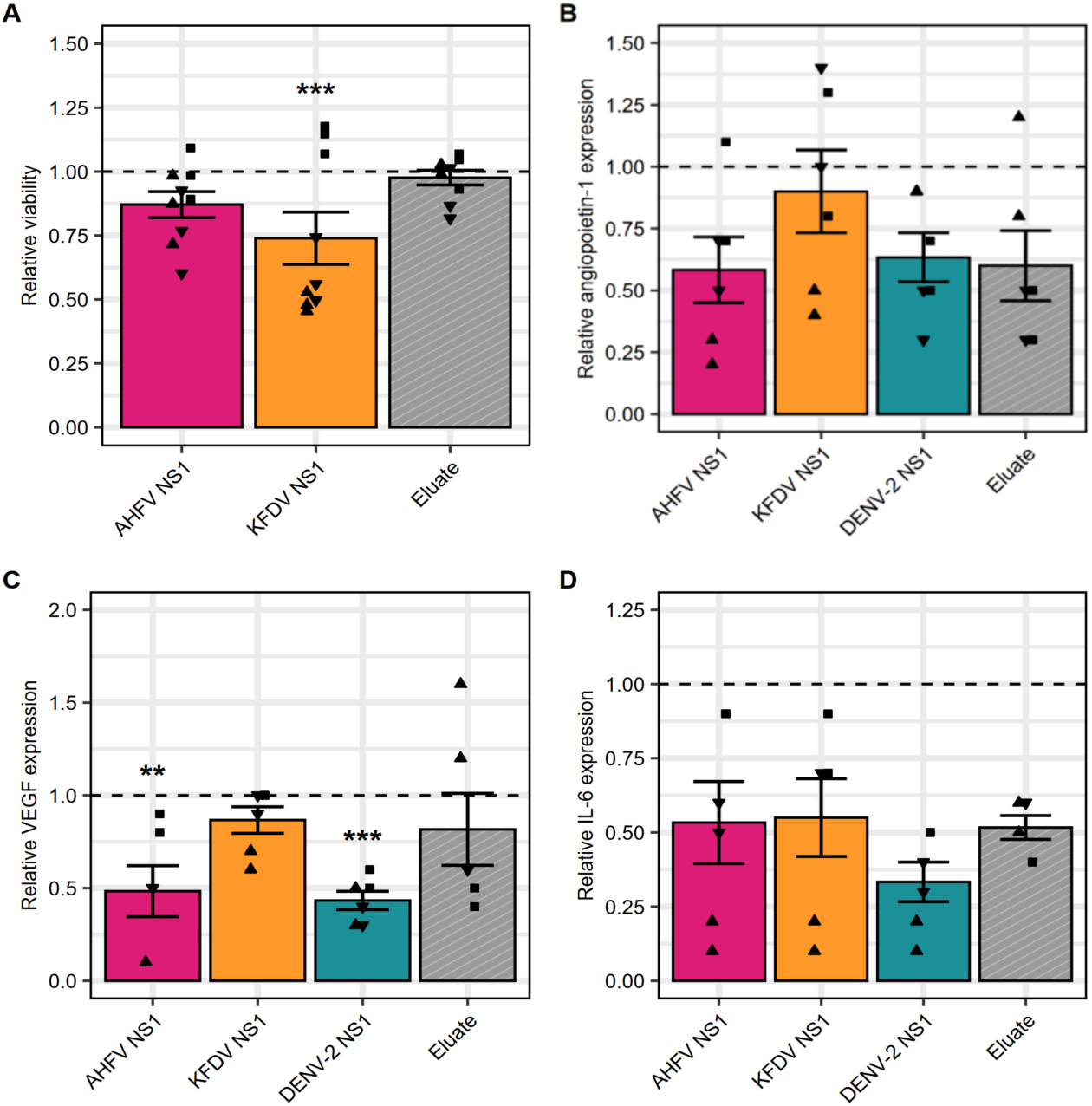
AHFV and KFDV NS1s differentially modulate pericyte physiology. (A) Viability of HSCs treated with 1 µg/mL NS1 for 24 h, relative to untreated HSCs (dashed line). (B–D) Relative expression of secreted factors known to influence endothelial function: angiopoietin-1 (B), vascular endothelial growth factor (VEGF) (C), and interleukin-6 (IL-6) (D) in HSCs following 24 h of NS1 treatment, normalised to untreated HSCs (dashed line). In all experiments, expression is shown relative to untreated HSCs, represented by a dashed line. Data points represent technical replicates within an experiment with different symbol shapes corresponding to different experiments (N=3, n=3 for the viability assays and N=3, n=2 for the RT-qPCR assays). Bars represent the mean, and error bars represent SEM. Statistical comparisons represent all pairwise differences between treatment groups. Indicated significance is for treatment versus untreated HSCs; * *P* < 0.05; ** *P* < 0.01; *** *P* < 0.001.

The expression of three secreted proteins known to influence microvascular function was assessed by RT-qPCR in HSCs following 24 h of NS1 treatment: angiopoietin-1, VEGF, and IL-6. AHFV NS1 significantly decreased VEGF expression, similarly to DENV-2 NS1 (Figure 3C), whereas AHFV NS1 did not significantly affect the expression of angiopoietin-1 or IL-6 (Figure 3B, D). In contrast, KFDV NS1 had no significant effect on the expression of any of the markers tested, suggesting that this protein may engage alternative signalling pathways compared to AHFV and DENV NS1. Taken together, these data show that both AHFV and KFDV NS1 modulate pericyte physiology in a number of ways that point to potential differences in the mechanisms by which these viruses affect the microvasculature.

### AHFV and KFDV NS1s do not induce hyperpermeability in pericyte-endothelial cell cocultures

To determine whether AHFV and KFDV NS1 were also able to induce hyperpermeability in pericyte-endothelial cell cocultures, we cultured HUVEC monolayers in the Electric Cell-substrate Electrical Impedance Sensing (ECIS) Z-Theta system, to measure the integrity of the endothelial barrier. We modelled the ability of NS1 to disrupt pericyte-to-endothelial cell paracrine signalling, which we previously demonstrated [9], by treating the endothelial cell monocultures with preconditioned medium from HSCs treated with NS1 for 24 h. Although media from TNF-α-treated HSCs increased permeability in HUVEC monolayers (Figure 4A), conditioned media from HSCs treated with AHFV or KFDV NS1 did not cause hyperpermeability compared to control media in either cell type over a 24 h time course. The same observations were made when preconditioned media from HSCs was added to liver sinusoidal endothelial cells (LSECs), a more representative model for liver microvascular permeability (Figure 4B).

**Figure 4:**
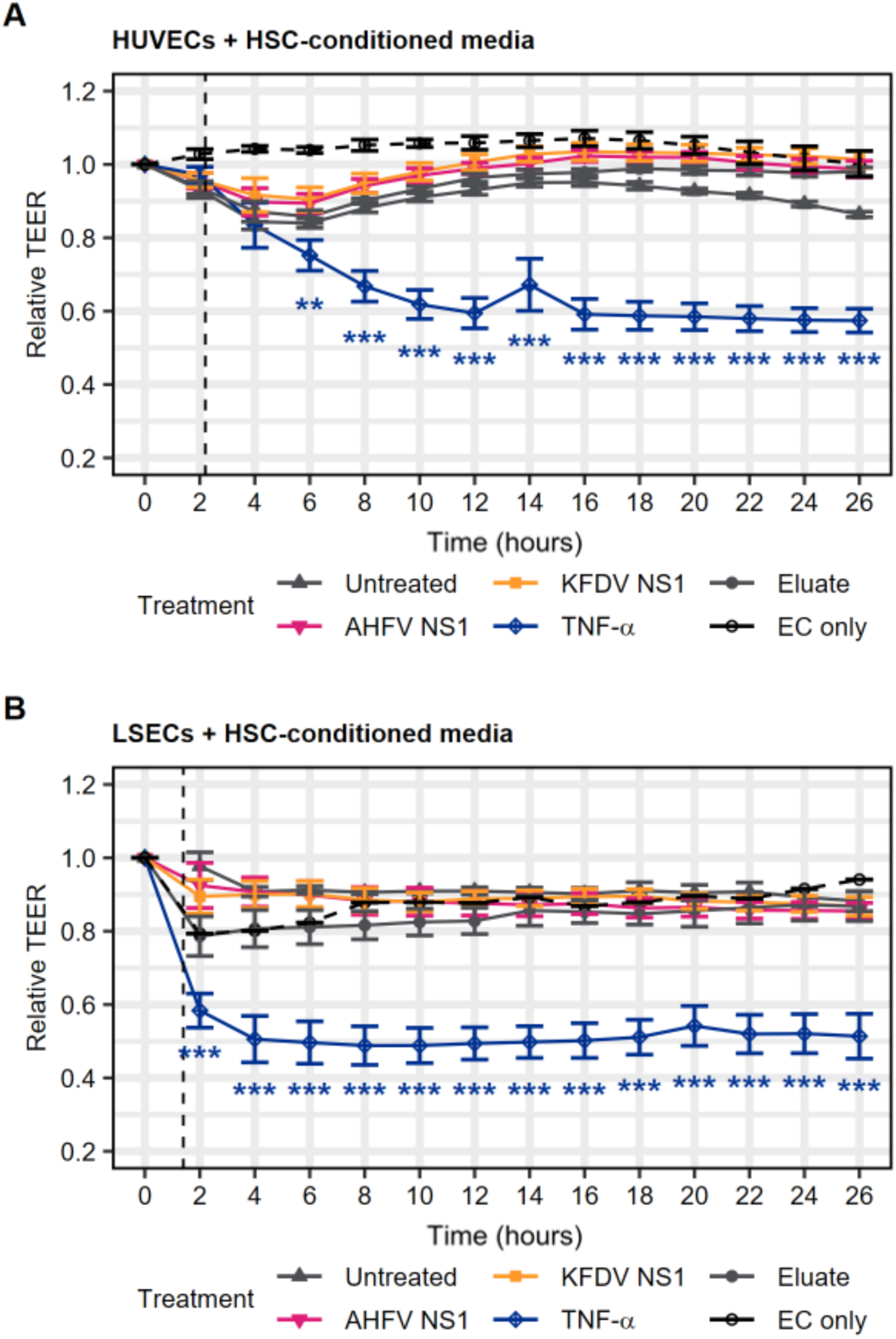
AHFV and KFDV NS1 do not induce hyperpermeability in endothelial monocultures following treatment with NS1-conditioned pericyte media. ECIS measurement of permeability of monocultures of HUVECs (A) or LSECs (B) treated with preconditioned media from HSCs treated with 1 µg/mL NS1 or 100 ng/mL TNF-α for 24 h; “Untreated” refers to HUVECs or LSECs treated with HSC media only and the “EC only” refers to untreated HUVECs or LSECs without the addition of HSC media. Each data point represents the mean **±** SEM (N=3, n=2). Statistical comparisons represent all pairwise differences between treatment groups at each time point. In all experiments indicated significance is compared to untreated cocultures; * *P* < 0.05; ** *P* < 0.01; *** *P* < 0.001.

Next, HUVECs or LSECs and HSCs were cultured on either side of a transwell insert in a semi-contact manner and the endothelial barrier permeability was measured by adding fluorescently-labelled dextran to the transwell insert 24 h after NS1 treatment and measuring its transfer across the endothelial barrier (Figure 5A-B). However, AHFV and KFDV NS1 had no impact on endothelial permeability in either coculture model (Figure 5A-B).

**Figure 5:**
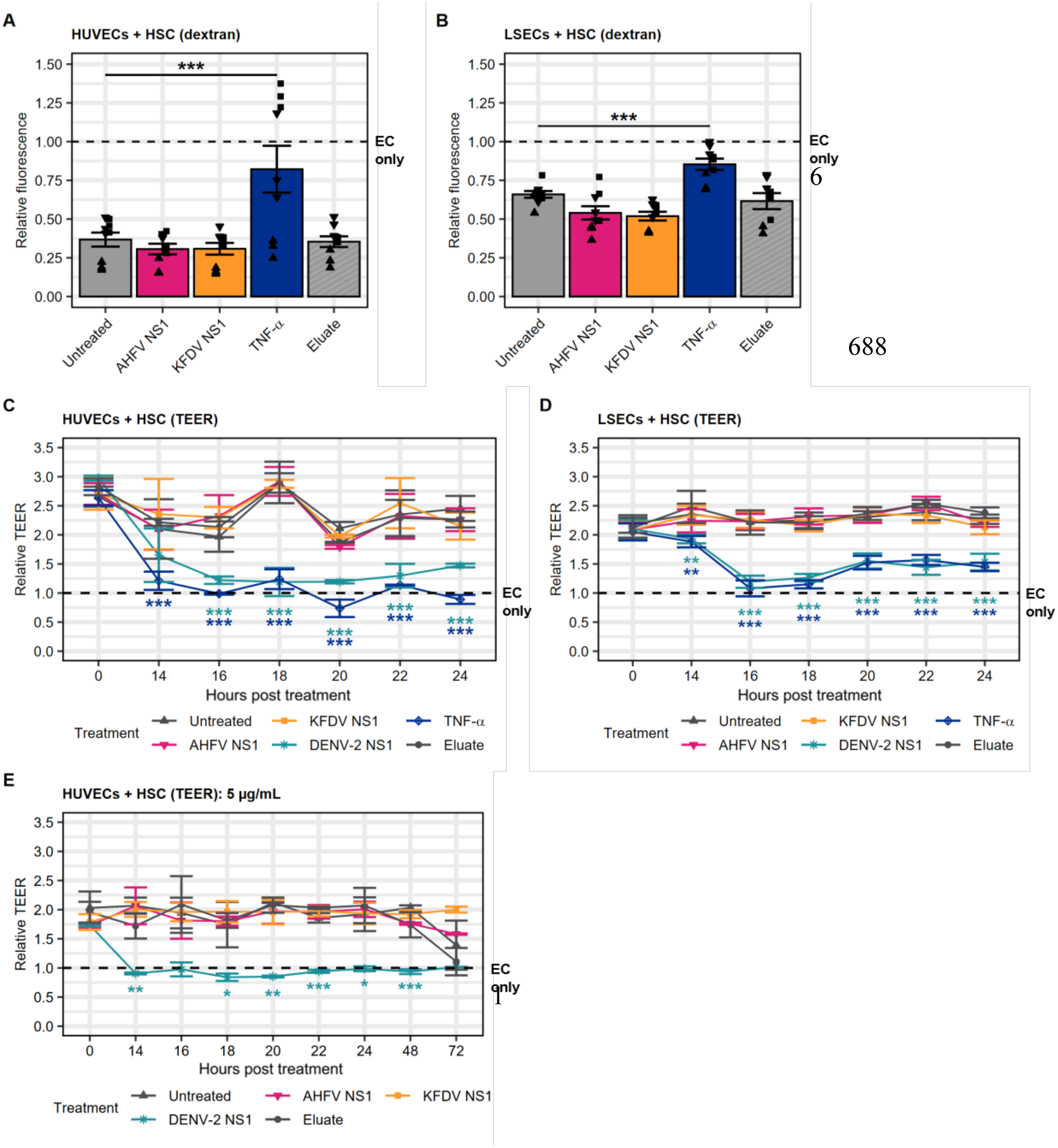
AHFV and KFDV NS1s do not induce hyperpermeability in pericyte endothelial cocultures. (A, B) Dextran permeability assay measuring HUVEC (A) or LSEC (B) permeability in coculture with HSCs 24 h post-treatment with 1 µg/mL NS1, 100 ng/mL TNF-α (positive control), or eluate (negative) control. Data points represent technical replicates within an experiment with different symbol shapes corresponding to different experiments (N=3, n=3). Bars represent the mean, and error bars represent SEM. (C-E) TEER measurement of semi-contact cocultures of HUVECs (C, E) or LSECs (D) cultured with HSCs and treated with 1 µg/mL NS1 (C-D) or 5 µg/mL NS1 (E), 100 ng/mL TNF-α (positive control), or eluate (negative) control for 24 h or HUVECs for 72 h (D). Permeability measurements were normalised to the untreated endothelial cell only control (EC only; dashed line), and each data point represents the mean **±** SEM (N=3, n=4 for panels A-C and N=1, n=3 for panel E). Statistical comparisons represent all pairwise differences between treatment groups at each time point. In all experiments indicated significance is compared to untreated cocultures; * *P* < 0.05; ** *P* < 0.01; *** *P* < 0.001.

We further confirmed our dextran results by measuring the transendothelial electrical resistance (TEER) using the EVOM as a measure of endothelial barrier integrity up to 24 hours post-treatment as before [9]. As expected, HSCs significantly increased the relative TEER (strengthened the endothelial barrier) by at least two-fold, and both DENV-2 NS1 and TNF-α significantly reduced this effect, completely abrogating the ability of pericytes to support the endothelial barrier (Figure 5C-D). However, in agreement with our dextran data, this experiment confirmed that AHFV and KFDV NS1 did not significantly increase hyperpermeability of the endothelial barrier compared to the controls when these proteins were tested at a concentration of 1 µg/ml. We also failed to see any impact of AHFV and KFDV NS1s at the higher concentration of 5 µg/ml up to 72 hours post-treatment, at which point the integrity of the endothelial barrier became compromised even in control conditions due to the extended culture period (Figure 5E). Overall, our data show that, unlike DENV-2 NS1, AHFV and KFDV NS1s do not induce hyperpermeability in several pericyte-endothelial cell coculture models, as measured through a range of permeability assays.

### AHFV and KFDV NS1s do not induce hyperpermeability in HUVEC or LSEC monocultures

Even in the absence of pericytes, NS1 from DENV and other orthoflaviviruses induces endothelial cell permeability, albeit to a lower degree than in the presence of pericytes [9,13,15,37]. Therefore, HUVECs were treated with DENV-2 NS1 and TNF-α (as a positive control), as well as with a DENV-2 N207Q NS1 mutant (negative control) defective in its ability to induce endothelial hyperpermeability [38]. As expected, DENV-2 NS1 and TNF-α significantly increased permeability in HUVEC monolayers by 14 h post-treatment, whereas the N207Q NS1 mutant had no effect at any timepoint (Figure 6A). In contrast, AHFV NS1 and KFDV NS1 failed to increase endothelial barrier permeability in both HUVEC and LSEC monocultures compared to the negative control (eluate), unlike the TNF-α positive control (Figure 6B-C). These data indicate that, similarly to our coculture model, AHFV and KFDV NS1 do not disrupt HUVEC or LSEC barrier integrity when endothelial cells are cultured in the absence of pericytes.

**Figure 6:**
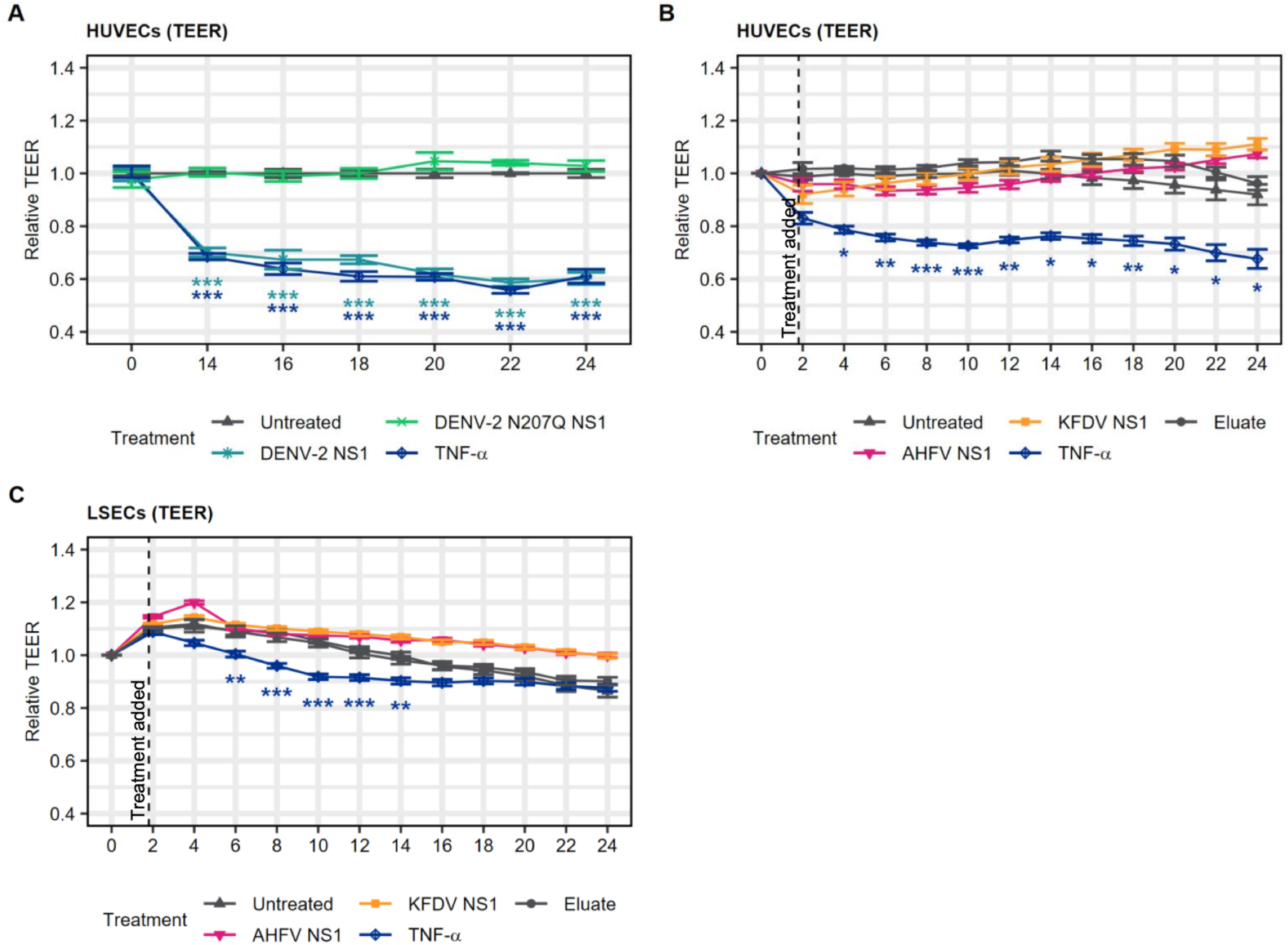
AHFV and KFDV NS1s do not induce hyperpermeability in HUVEC or LSEC monocultures. Endothelial barrier integrity measurements for HUVEC (A, B) or LSEC (C) monocultures following treatment with 1 µg/mL wild-type DENV-2, AHFV or KFDV NS1, or the nonfunctional DENV-2 NS207Q mutant (negative control), or 100 ng/mL TNF-α (positive control), or eluate (negative) control as indicated. Permeability measurements were normalised to the untreated endothelial cell only control (Untreated) for EVOM TEER data (A), or to each sample before treatment for ECIS data (B, C). Each data point represents the mean **±** SEM (N=3, n=2 for panel A and N=3, n=2 for panels B-C). Statistical comparisons represent all pairwise differences between treatment groups at each time point. Indicated significance is for treatment versus untreated EC only control (Untreated); * *P* < 0.05; ** *P* < 0.01; *** *P* < 0.001.

## Discussion

In this study, we showed that AHFV and KFDV NS1 disrupt the ability of pericytes to physically interact with and functionally support endothelial cells in a 3D angiogenesis assay. AHFV NS1 additionally altered pericyte signalling pathways, including VEGF signalling, while KFDV reduced HSC viability. Taken together, these findings show that both AHFV and KFDV NS1 modulate pericyte physiology in a number of different ways, but do not affect endothelial barrier integrity in pericyte-endothelial cell cocultures or when HUVECs or LSECs were cultured alone, at least not with the endothelial cells tested here. These data differ from DENV-2 NS1, which disrupts the ability of pericytes to support endothelial cell function in both angiogenesis and permeability assays [9]. We therefore propose that NS1 may need to cause dysregulation of pericytes in combination with dysregulation of endothelial cells to induce maximum levels of microvascular permeability, although this needs to be conclusively demonstrated.

Our data differ from a wide range of orthoflavivirus NS1s that impair endothelial barrier integrity in the absence of pericytes [11,12,15,37,39]. It should be noted that NS1-induced permeability has been shown to be tissue-specific. For example, neurotropic orthoflaviviruses such as WNV and Japanese encephalitis virus only affect brain endothelial cells, ZIKV affects brain and placental endothelial cells, whereas DENV, which causes systemic disease, affects endothelial cells from multiple different tissues [37]. Recent studies indicate that this tissue specificity also applies to tick-borne flaviviruses. Tick-borne encephalitis virus (TBEV), for example, did not trigger hyperpermeability in HUVEC monocultures but significantly increased permeability in lung endothelial cell monocultures [39]. In our study, we selected HUVECs because they are a widely used system and chose liver microvascular cells due to reported elevated liver enzyme levels in AHFV- and KFDV-infected patients, as well as liver pathology in animal models [4,6–8,35]. We cannot exclude the possibility that AHFV and KFDV NS1 may dysregulate endothelial barrier integrity in endothelial cells derived from other organs. The absence of an observable response in the two endothelial cell types and under the experimental conditions examined here does not preclude activity in other vascular beds and contexts. This is particularly relevant given that patients present with a range of symptoms, including gastrointestinal and neurological manifestations [4,6–8]. Furthermore, the concentration of NS1 in patient serum is unknown for AHFV and KFDV; our experiments used NS1 concentrations based on DENV data [40], and it is possible that higher levels of AHFV or KFDV NS1 are required to induce endothelial permeability. We also note that the DENV-2 NS1 used in this study was commercially sourced, whereas the AHFV and KFDV NS1 proteins were produced in-house. While all proteins were handled and tested under comparable conditions, we cannot exclude the possibility that differences in expression or purification pipelines may influence protein activity.

Nevertheless, our data potentially untangle the importance of direct endothelial cell effects in the context of NS1-induced hyperpermeability. While the permeability and angiogenesis assays we used assess different aspects of microvascular cell functional interactions, we previously observed that DENV-2 NS1-induced effects in one model translated to the other [9]. Interestingly, as VEGF is a key regulator of angiogenesis, controlling endothelial activation, migration, and vessel formation, and contributing to pericyte–endothelial communication [41–43], the modulation of VEGF by AHFV NS1 would be expected to impair endothelial barrier integrity. Furthermore, the reduced viability of HSCs observed after KFDV NS1 treatment might be anticipated to reduce the ability of HSCs to support the endothelial barrier. The fact that AHFV and KFDV modulate pericyte physiology in multiple ways, and yet endothelial permeability is not observed here in the absence of impacts on endothelial cells themselves is suggestive of endothelial cell dysfunction potentially being required for orthoflavivirus microvascular leakage. Our data also reveal potentially interesting differences in how these tick-borne orthoflaviviruses modulate pericyte function that should be further explored in future studies.

As well as further investigation into the effects AHFV and KFDV NS1s have on a range of endothelial cells and pericytes from different tissues, future work should also focus on elucidating the mechanisms underlying the lack of permeability changes observed in the present models. In particular, identifying NS1 receptors would enable investigation of binding and internalisation kinetics, which may help explain the lack of barrier disruption in these cell types. Previous studies have shown that disruption of NS1 glycosylation sites can abrogate its internalisation and associated permeability effects [38]. Although sequence analysis in this study did not identify mutations at these sites, other factors may still limit NS1 binding or uptake, warranting further investigation.

Here, we conclude that while pericytes are important for amplifying the impact of NS1 on endothelial barrier integrity, disruption of pericyte function alone may be insufficient to induce NS1-mediated microvascular hyperpermeability in the absence of direct effects on endothelial cells. Our data highlight a need to gain a better understanding of the mechanisms by which orthoflavivirus NS1s disrupt pericyte-endothelial cell interactions, and the relative contributions each cell type makes to NS1-induced microvascular leakage.

## Author Contributions

Conceptualization: PC, KM

Data curation: EB, JD

Formal analysis: EB, JD

Funding acquisition: PC, KM

Investigation: EB, JD

Methodology: EB, JD, PC, KM

Supervision: PC, KM

Visualization: EB

Writing – original draft: EB

Writing – review & editing: all authors

## Acknowledgements

We thank Eva Harris (University of California, Berkeley, CA USA) for allowing the use of the DENV-2 NS1 N207Q mutant. We also thank Simon Gubbins (The Pirbright Institute) and Giovanni Lo Iacono (University of Surrey) for their continued supervision and support of EB.

